# Data mining reveals the diversity of prophage endolysins targeting pathogenic enterococci

**DOI:** 10.64898/2026.04.26.720912

**Authors:** Finn O’Dea, Andrew Kinsella, Brooks J Rady, Andrew D Millard, Joseph Brown, Graham P Stafford, Stéphane Mesnage

## Abstract

Antimicrobial resistance (AMR) poses a critical global health threat, with enterococci among the leading contributors due to their intrinsic and acquired resistance to antibiotics. Clinically relevant species, including *Enterococcus faecalis* and *Enterococcus faecium*, as well as the emerging poultry pathogen *Enterococcus cecorum*, highlight the need for alternative therapeutics across human and agricultural settings. Bacteriophages and their derived enzymes, particularly endolysins, offer promising antibacterial strategies but challenges such as phage resistance and limited lysin diversity hinder their application. In this study, we performed a large-scale analysis of prophage-encoded endolysins across these three enterococcal opportunistic pathogens, characterizing over 48,000 sequences. We identified 33 distinct domain architectures combining diverse catalytic and cell wall-binding domains, including novel putative cell wall binding domains. These findings expand the known diversity of enterococcal lysins and provide a comprehensive resource for the rational design of stable, recombinant “enzybiotics” to combat multidrug-resistant enterococcal infections.

**Data summary:** All genomes analysed in this work are available through Genbank. The data mining strategy was carried out open-access software available through GitHub as described in the Methods section. The raw output of the search and sequences obtained after each filtering step are provided in Supplementary Files 1 and 2. Modelling data related to figure 6 is provided in supplementary File 3.

**Impact statement:** Antimicrobial resistant enterococci threaten therapeutic options in both medicine and agriculture. Yet, the therapeutic potential of bacteriophage-derived endolysins (enzybiotics) is limited by an incomplete understanding of their natural diversity. By analysing more than 48,000 prophage encoded lysins from *E. faecalis, E. faecium*, and *E. cecorum*, this study provides the most extensive characterization of enterococcal lysin architectures to date. The identification of 34 distinct domain organizations, including a previously unrecognized cell wall–binding domain, substantially broadens the known functional repertoire of these enzymes. This work fills a major knowledge gap and offers a foundational resource for engineering stable, targeted enzybiotics to combat multidrug resistant enterococcal infections.

## Introduction

Antimicrobial resistance (AMR) is a growing global health crisis, projected to result in approximately 10-40 million deaths annually by 2050 if current trends continue (Naddaf, 2024). Among the major contributors to the global antimicrobial resistance (AMR) crisis are enterococci. Enterococcal infections are especially prevalent in healthcare settings and are associated with high morbidity and mortality due to their intrinsic resistance to multiple antibiotics and their capacity to acquire additional resistance genes (Hollenbeck and Rice, 2012, Kristich et al., 2014). Enterococci are opportunistic pathogens responsible for a broad range of infections, including bloodstream and urinary tract infections (Codelia-Anjum et al., 2023), endocarditis (Beganovic et al., 2018), and device-associated infections (Donelli and Guaglianone, 2004). *Enterococcus faecalis* (*Efs*) and *Enterococcus faecium* (*Efm*) are the two species of greatest clinical relevance, with vancomycin-resistant *E. faecium* (VRE) being designated a high-priority pathogen by the World Health Organization (Organization, 2014; Health and Services, 2019). This classification reflects both its rising clinical significance and the substantial challenges in treating it via traditional antimicrobial therapy.

Beyond their threat to human health, enterococci are increasingly recognized as pathogens in agricultural settings, particularly in poultry. Over the past decades, the species *Enterococcus cecorum* (*Ecm*) has emerged as a prominent poultry pathogen, accounting for up to 7% of infections in the broiler farms of some countries (Souillard et al., 2022). Antimicrobial resistance has been cited as a significant contributor to the rise in these infections (O’Dea et al., 2025, Laurentie et al., 2023, Dolka et al., 2016). The emergence of antibiotic-resistant enterococci in both human and agricultural populations underscores the need for a novel class of therapeutics, particularly given the declining pace of antibiotic development for these pathogens. *Efs, Efm* and *Ecm* have diverged from each other by a long evolutionary history and have been assigned to three distinct, deeply-branching, clades within the genus *Enterococcus* (*Efs*: clade I, *Efm*: clade II, *Ecm*: clade IV) (Schwartzman et al., 2023).

Bacteriophages (phages) have emerged as a promising alternative or adjunct to antibiotics (Hatfull et al., 2022) with multiple reports demonstrating successful phage therapy in clinical and compassionate-use settings (Dedrick et al., 2022, Onallah et al., 2023). However, bacterial resistance to phages can arise rapidly through a variety of mechanisms, including the modification of surface receptors, activation of restriction-modification systems, and other phage defence strategies (Teklemariam et al., 2023). Such resistance limits the long-term effectiveness of phage therapy when used alone.

A complementary approach that could help overcome phage resistance involves the use of enzymes that cleave bacterial peptidoglycans (the essential component of the cell envelope), thereby inducing lysis. These enzymes called “enzybiotics” were originally defined as antibacterial proteins with enzymatic activity (Nelson et al., 2001). Enzybiotics are most commonly phage endolysins that degrade the peptidoglycan at the end of the infection cycle to release virions (Gondil et al., 2020, Rodríguez-Rubio et al., 2016). Although phage tail-associated proteins also contain peptidoglycan-hydrolytic domains (Alrafaie and Stafford, 2023), their therapeutic potential has been minimally explored, with the notable exception of the *Efs* metallopeptidase EnpA (Mitkowski et al., 2024, Feldwisch et al., 2010). Unlike whole phage, endolysins do not require adsorption to a specific receptor or intracellular replication to exert their bactericidal effect. Additionally, endolysins attack peptidoglycan, thereby reducing the likelihood of resistance being developed (Loeffler et al., 2001). Multiple studies have demonstrated the efficacy of endolysins against diverse enterococcal strains, including VRE, with reports of bacterial resistance remaining exceedingly rare (Kim et al., 2020, Fischetti, 2005). Despite this promise, the translation of endolysins to clinical application has been limited by challenges in scaling up production, including poor protein solubility and stability, and a relatively limited pool of natural lysin candidates to draw from during therapeutic development (Liu et al., 2023, Balaban et al., 2022). Recent studies have investigated the diversity of tail-associated lysins (Alrafaie and Stafford, 2023) and endolysins encoded by 171 virulent phages targeting enterococci (Viglasky et al., 2023), providing a useful resource to explore the therapeutic activity of these enzymes.

Building upon this research, our work presents a large-scale analysis of prophage-encoded endolysins within *Efm, Efs* and *Ecm*. Using a high-throughput pipeline, we characterized over 48,000 endolysin sequences and identified 33 unique domain architectures, modularly combining diverse catalytic (CD) and cell-wall-binding (CWBD) domains. We selected distinct amino acid sequences for each domain architecture that could be used to produce recombinant enzymes with therapeutic potential. Our analysis revealed a diverse repertoire of enterococcal phage lysins spanning multiple enzymatic classes and associated with a wide range of CWBDs, including new putative binding domains. This work establishes a foundational resource for the rational design of novel endolysin-based therapeutics aimed at combating multidrug-resistant enterococci in both clinical and agricultural settings.

## Materials and Methods

### Prophage Prediction

Genomes were downloaded from NCBI using Biopython v1.81 Bio.Entrez module, based on a list of all genomes identified as *Enterococcus* (taxid: 1350) that were present on Genbank at time of download (December 2023). Any genomes with <90% completeness or >10% contamination according to CheckM v1.2.3 (Parks et al., 2015) using ‘lineage_wf’ were discarded, and GTDB-Tk v2.4.0 (Chaumeil et al., 2022) using ‘classify_wf’ was used to determine bacterial species. Prophages were identified with PhageBoost v0.1.7 (Sirén et al., 2021), then run through VIBRANT v1.2.1 (Kieft et al., 2020) and geNomad v1.5.2 (Camargo et al., 2024) to check, with only prophages confirmed by PhageBoost and either VIBRANT or geNomad being kept. If both VIBRANT and geNomad identified it as a prophage, the longer genome was kept (the VIBRANT prediction was kept if both were the same length). Prokka v1.14.6 was used to annotate genomes using the PHROGS database (Terzian et al., 2021).

### Identification of prophage-encoded endolysins and selection of representative endolysins

PHROGS annotations were used to identify genes annotated as “endolysin”. “Hypothetical proteins” encoded by genes one or two genes away from a holin were also selected as putative endolysins. From the raw output of the search, identical sequences along with sequences less than 115 amino acids (deemed to not encode complete domains) were removed. Following this step, Clustal Omega (Sievers et al., 2011) was used to identify and remove sequences which displayed more than 95% sequence identity. Interpro (Blum et al., 2025) and the NCBI conserved domain database (Wang et al., 2023) were then used to identify domains and to remove any truncated proteins which only made up a portion of an intact catalytic domain. AlphaFold (Abramson et al., 2024) predictions were performed to detect putative binding domains not detected by Interpro searches. Putative binding domains were subsequently run through Foldseek (van Kempen et al., 2024) to identify the closest homologues. BioEdit was used to generate sequence identity matrices and to align those sequences.

### *In silico* docking of enterococcal muropeptides into putative binding domains

Muropeptides from *Efs* (gmgm-AQK[AA]AA∼gm-AQK[AA]AA∼g) and *Efm* (gmgm-AQK[D]AA∼gm-AQK[D]AA∼g) were docked into the putative cell wall binding domains BSD, CTD1, CTD2, and DUF5648 using Boltz v2.2.1 (Passaro et al., 2025). To differentiate between genuine and spurious docking results, dockings with several positive control domains (LysM, SH3) and negative control proteins (GFP, RNBR) were also predicted (see File S3, ‘Binding Analysis/Pluto Notebook.jl’ or https://pluto.land/n/s8dxd3y8 for a complete list of amino acid sequences). SMILES structures for each muropeptide were computed using PGFinder (Alamán-Zárate et al., 2025), and manually unreduced (by default, all SMILES structures generated by PGFinder contain reduced terminal sugars). Each SMILES structure was then paired with an amino acid sequence in a YAML file and fed to ‘boltz predict’ alongside the following flags: ‘--use_msa_server --use_potentials -- step_scale 1 --recycling_steps 10 --diffusion_samples 25’. The full set of input YAML files and Boltz outputs can be found in File S3 under ‘Full Boltz-2 Results/’.

The ‘ligand_iptm’ scores for each protein-muropeptide docking were extracted using a custom Pluto.jl notebook (Bezanson et al., 2017, Giazitzidou, 2018) and visualized as a box plot using Makie.jl (Danisch et al., 2021). The custom “Docking Consistency” metric was calculated in two steps: first, ligand atoms contacting the protein (< 8□) in at least 13 of the 25 Boltz predictions were selected using BioStructures.jl (Greener et al., 2020), and ligand atoms consistently contacting at least 10 protein atoms were determined to be “docked atoms” (this selects for binding pockets or grooves as opposed to tangential points of contact). The final “docking consistency score” was then calculated as the number of docked atoms divided by their RMSD when compared to the top-ranked Boltz model. The full Pluto.jl Notebook code can be found in File S3, ‘Binding Analysis/Pluto Notebook.jl’, or at https://pluto.land/n/s8dxd3y8. Finally, hydrogen bonding between the most confident model of BSD and the *Efs* muropeptide was computed and visualized in ChimeraX (Pettersen et al., 2021).

## Results and discussion

### Identification of putative endolysins encoded by prophages in pathogenic enterococci

We sought to leverage the vast collection of available prophage genome sequences as an untapped resource for the discovery of novel enzybiotics. A total of 29,592 enterococcal genomes were analysed leading to the identification of 48,399 putative prophage endolysins across the three species chosen (Supp File 1): 299, 21989 and 26111 sequences for *Ecm, Efm* and *Efs*, respectively. After removing identical sequences and retaining only those longer than 115 residues (unlikely to contain any of the catalytic domains of interest) approximately 10,000 candidate sequences remained. A final selection step retaining only endolysins with less than 95% sequence identity and those annotated to have enzymatic domains resulted in a refined dataset containing 245 endolysins across the three species (37 in *Ecm*, 82 in *Efm* and 126 in *Efs*) (Table 1).

**Table 1.** Search strategy for endolysin identification. The output of each search step is provided in Supp. File 1; All sequences resulting from the Interpro searches are in Supp. File 2.

| Search step | Number of sequences |  |  |
| --- | --- | --- | --- |
|  | *Ecm* | *Efm* | *Efs* |
| Raw output | 299 | 21,989 | 26,111 |
| Remove redundant sequences | 138 | 4,177 | 6,283 |
| Remove size (>115 residues) | 106 | 3,845 | 6,103 |
| Clustal Omega (<95% identity) | 65 | 289 | 373 |
| Interpro (truncated domains) | 37 | 82 | 126 |

Endolysins can be broadly classified into four groups based on the peptidoglycan (PG) bond they cleave (Fig. 1). The first class corresponds to glycosyl hydrolases that target the glycan chain and includes *N*-acetylmuramidases and *N*-acetylglucosaminidases. Both cleave the β-1,4-glycosidic bonds between *N*-acetylmuramic acid (MurNAc) and *N*-acetylglucosamine (GlcNAc) residues but muramidases generate a MurNAc reducing end whilst glucosaminidases release a GlcNAc reducing end. The second class corresponds to endopeptidases, which cleave peptide bonds either in the pentapeptide stem or in the lateral chain linking adjacent stem peptides. Endopeptidases can be stereospecific, cleaving either L,D or D,L bonds. The third class corresponds to *N*-acetylmuramoyl-L-alanine amidases (amidases), which cleave the bond between the first L-alanine in the peptide stem and MurNAc.

**Figure 1.**
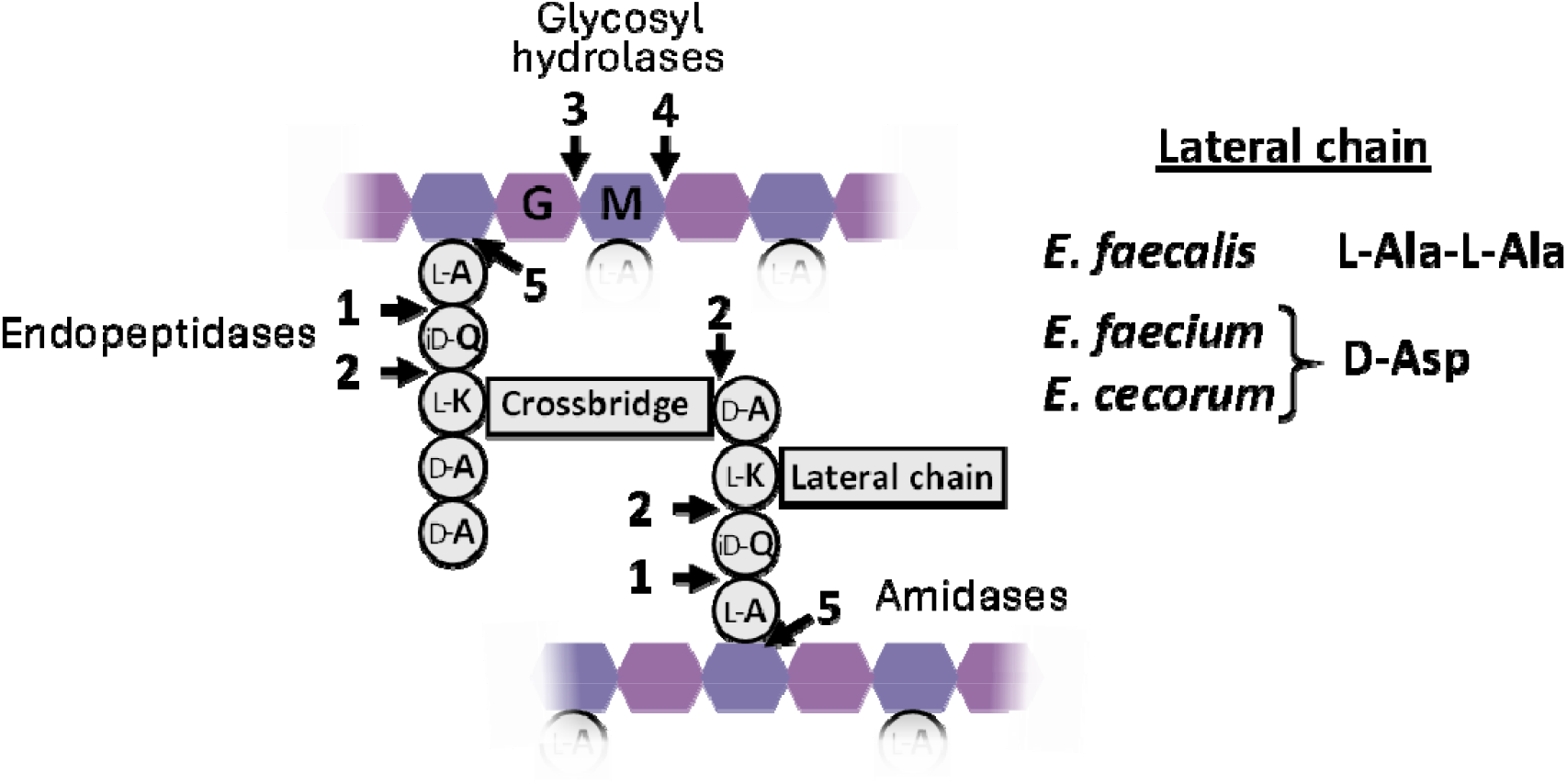
Peptidoglycan structure of selected enterococcal species and bonds cleaved by endolysins. Arrows indicate cleavage sites for different classes of enzymes. 1, L,D endopeptidases; 2, D,L endopeptidases; 3, *N*-acetylglucosaminidases (generate glycan chains with a GlcNAc at their reducing end); 4, *N*-acetylmuramidases (generate glycan chains with a MurNAc at their reducing end); 5, *N*-acetylmuramoyl-L-alanine amidases.

Catalytic domains (CD) belonging to the four classes of endolysins described above were usually found associated with cell wall binding domains (CWBDs) located at the C-terminus. A limited number of CDs were not associated with any CWBD. Interestingly, some endolysins displayed a more complex domain organisation, combining several CD and/or CWBDs.

### Endolysins with endopeptidase domains

Analysis of enterococcal prophage genomes identified endolysins with four distinct endopeptidase domains: NlpC/P60 (PF0087), CHAP (Cysteine, Histidine-dependent Amidohydrolases/Peptidases; PF00527), Amidase_5 (PF05382), and M23 metallopeptidases (PF01551). M23 domains were only found in *Efm* prophage-encoded endolysins whilst the three other catalytic domains were present in all species studied. All these domains were found associated with other CDs (see later section), but NlpC/P60, CHAP, and Amidase_5 domains were also found in endolysins with a single CD (Fig. 2). Species-specific patterns of domain organisation were found. Lysins encoded by *Efs* contained NlpC_P60, CHAP, and Amidase_5 as standalone endopeptidase CDs, whereas those identified in *Efm* and *Ecm* harboured NlpC/P60 or Amidase_5 domains, suggesting a lineage-specific diversification of phage-encoded lytic systems. CHAP and NlpC/P70 endolysins are well characterized and exhibit potent antimicrobial activity against enterococci and other Gram-positive bacteria (see Fischetti, 2005, and Loessner, 2005). For example, the CHAP-domain-containing lysin LysEF-P10 displays strong lytic activity against *Efs* strains (Cheng et al., 2017), and the NlpC/P60-containing LysPEF1-1 exhibited efficient bacteriolytic activity against multidrug resistant *Efs* and *Efm* strains (Wang et al., 2024). The Amidase_5 domain is most likely to display endopeptidase activity rather than amidase activity since it is structurally related to peptidases with a papain-like fold. No experimental evidence has shown the exact bond cleaved by the Amidase_5 CD, but it has been suggested that it could have D-Glutamine-L-Lysine endopeptidase activity (Kovalskaya et al., 2019).

**Figure 2.**
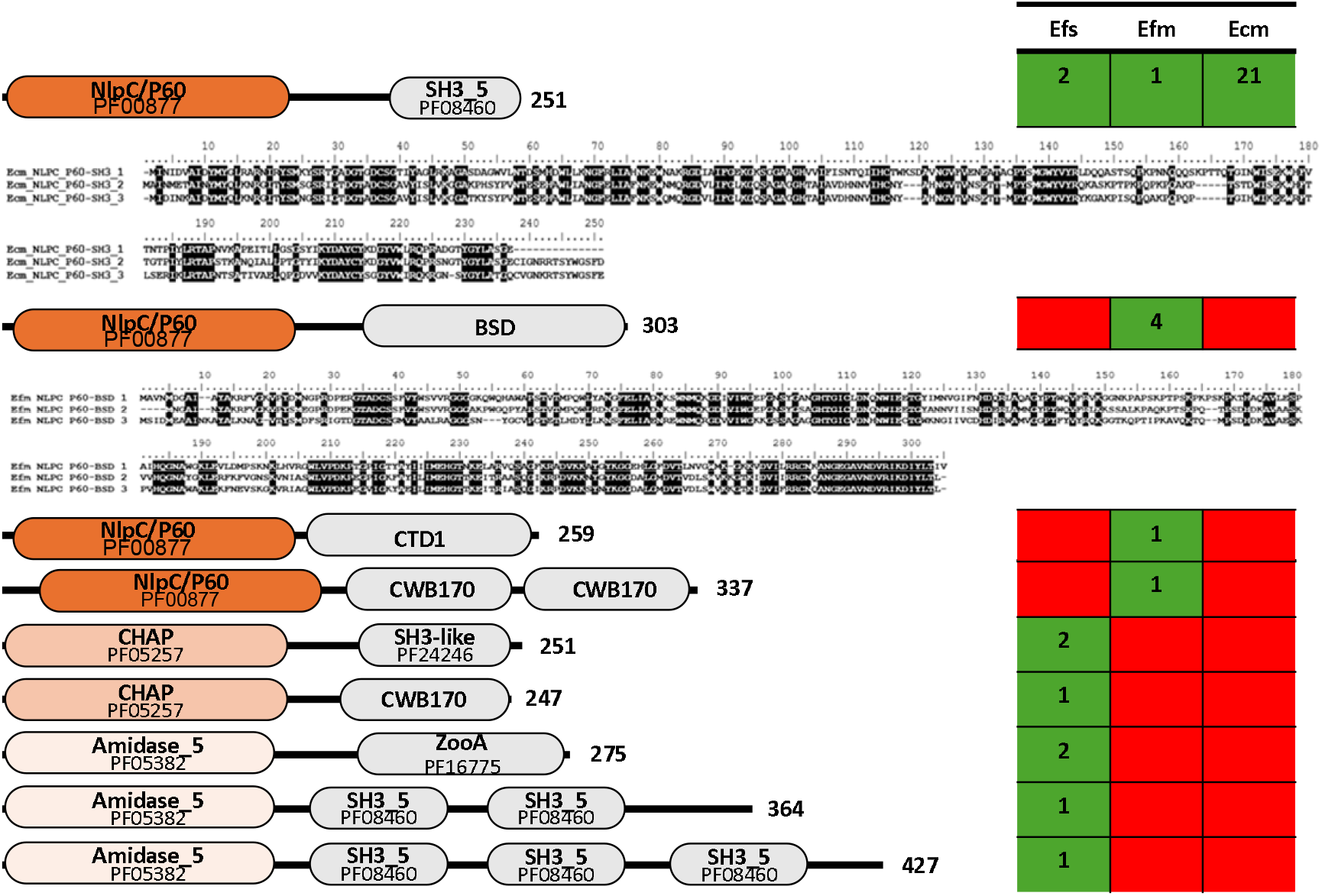
Domain architecture of endolysins containing a single endopeptidase CD. Domain organisations of endopeptidases combined with various CWBDs are shown. Boxes on the right-hand side show the number of sequence clusters (95% identity) in each species with a given domain architecture. An arbitrary cutoff of 75% identity led to the identification of three divergent groups of sequences for NlpC/P60-SH3 and NlpC/P60-BSD endolysins.

A total of nine distinct domain architectures combining a single endopeptidase CD with one or two putative CWBDs were identified. CWBDs included SH3 (SRC homology 3) domains (PF08460 or PF24246), CWB170 domains (PDB 6L00) or ZooA (PF16775), described in the literature (Gonzalez-Delgado et al., 2020, Xu et al., 2021, Lai et al., 2002). Two other uncharacterized putative CWBD were identified and named the beta-sandwich domain (BSD) based on its predicted structure and the C-terminal Domain 1 (CTD1). These domains were distributed differently between species and contributed to the observed diversity in lysin organization. CWB170, BSD or CTD1 were not detected by Interpro and were identified using AlphaFold and/or Foldseek.

Structural prediction and alignment of the CDs revealed a high degree of conservation in overall fold architecture. Pairwise structural alignments of CDs (e.g., NlpC/P60 or CHAP) from distinct species yielded TM-scores exceeding 0.7 across all comparisons, indicating significant structural similarity despite sequence divergence. Notably, catalytic domains from *Efm* and *Ecm* displayed greater structural similarity to one another than to those from *Efs*, consistent with their similar peptidoglycan composition. We further investigated amino acid sequence diversity within specific domain organizations. When a more stringent cutoff of 75% identity was set, we identified three divergent groups of sequences for two of the endopeptidase domain organisations described in Fig. 2 (NlpC/P60-SH3 and NlpC/P60-BSD) (Supp. File 2).

### Endolysins with amidase domains

Endolysins displaying distinct types of amidase CD were identified (Fig. 3). These enzymes belonged to the Amidase_2 (PF01510) superfamily (which includes peptidoglycan recognition proteins (PGRPs) and CwlA-like autolysins), the Amidase_3 (PF01520) superfamily, and one domain in the Amidase/PGRP superfamily (IPR036505). Amidase-containing lysins have been widely reported as effective antimicrobials against enterococci. Amidases such as PlyV12 or ORF9 exhibit broad lytic activity against vancomycin-resistant enterococci and display bactericidal activity against both *Efm* and *Efs* (Uchiyama et al., 2011, Yoong et al., 2004). Only one amidase CD was found in multi-catalytic enzymes; these CDs were mostly found in combination with a large variety of CWBDs, including SH3, CWB170, and the previously described BSD and CTD1 domains, mirroring the modular architecture observed for endopeptidase lysins. Amidase endolysins were also found in association with ZooA binding modules and CW_7 CWBDs (PF08230; Bustamante et al., 2017), domains which were not observed in endopeptidase lysins. Additionally, more complex CWBD arrangements were identified alongside the amidase domains, with multiple SH3 domains belonging to either the same or different classes being found at the C-terminus of amidase CDs. Species-specific patterns of catalytic domain distribution were again evident. Amidase_3 domains were detected exclusively in *Efm*, whereas Amidase_2 domains were distributed across all three species examined (*Efs, Efm*, and *Ecm*). Two of the three CWBDs identified in amidase-containing lysins (BSD and SH3) were present in all three species, while CWB170 was restricted to *Ecm* and *Efs*. All pairwise structural comparisons of identical amidase CDs yielded TM-scores exceeding 0.7, indicating a high degree of fold similarity across species. TM-scores among amidase domains were more uniform than those observed for endopeptidase domains, likely reflecting the conserved biochemical target of these enzymes, which are all predicted to cleave the same MurNAc-L-Ala bond in peptidoglycan. High sequence diversity was observed in amidase domain organisations. Four of the amidase-lysin domain architectures contained sequence clusters with less than 75% sequence identity. These included those encoding Ami2-BSD (10 *Efm* clusters), Ami2-SH3-SH3 (4 *Efs* clusters; 2 *Efm* clusters; 3 *Ecm* clusters), Ami2-CWB170 (2 *Efs* clusters) and Ami3-BSD (4 *Efm* clusters) architectures. Representative cluster sequences are provided in (Supp. File 2).

**Figure 3.**
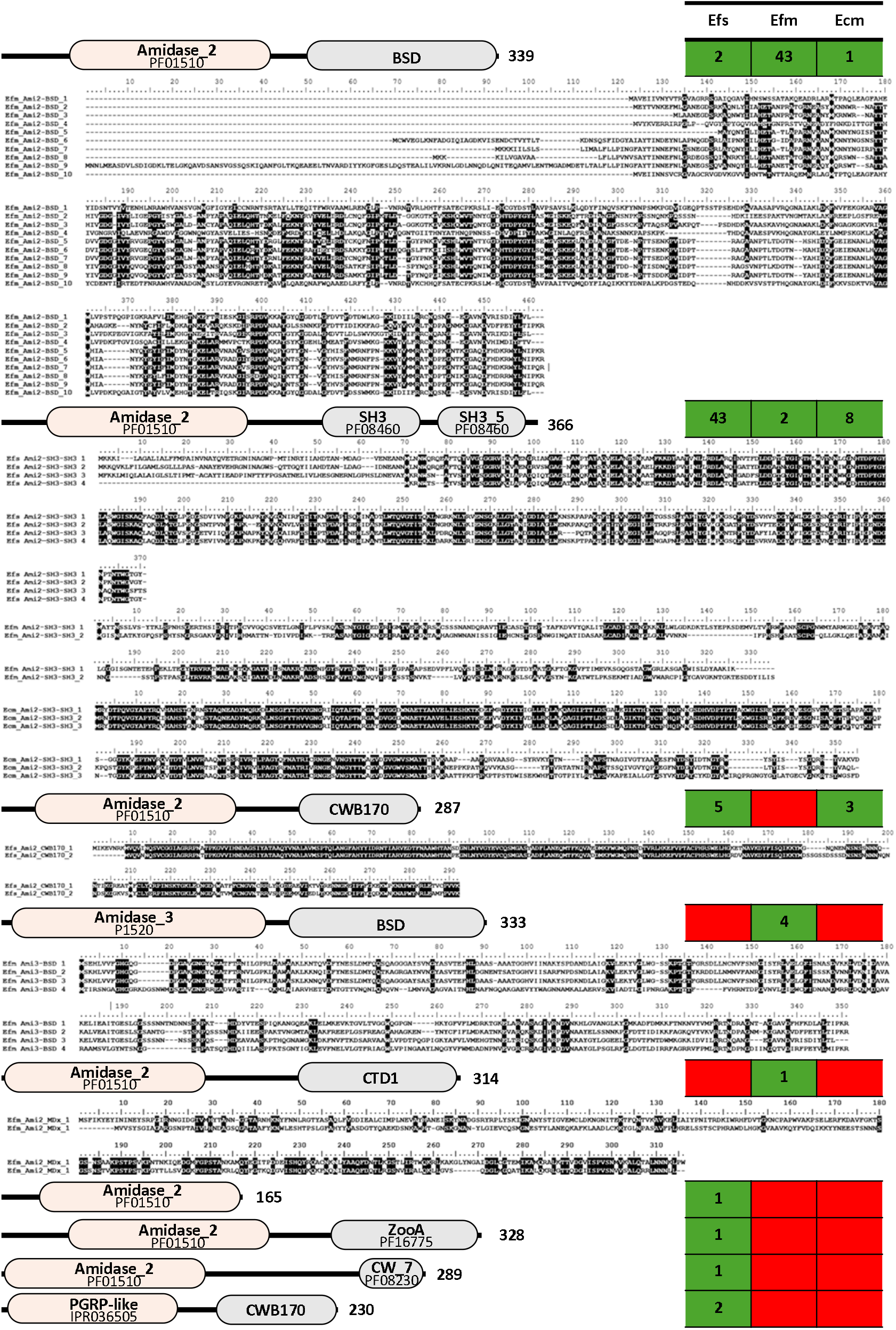
Domain architecture of endolysins containing an amidase CD. Domain organisations of multiple amidase CDs combined with various CWBDs are shown. Boxes on the right-hand side show the number of sequence clusters (95% identity) in each species with a given domain architecture. Further clustering at 75% identity led to the identification of divergent groups of sequences for several amidase endolysins.

### Endolysins with predicted muramidase domains

A relatively large number of endolysins contained a CD with muramidase activity. Three distinct CDs with this activity were identified: GH25 (PF01183), phage lysozyme (PF00959), and Lyz-like (PF13702), with GH25 being the most common (Fig. 4). Only one muramidase domain was found associated with another CD, which contrasted with the endopeptidase domains which were often found in combination with other CDs. Few examples of endolysins with muramidase activity have been described in the literature, except for *Efs* AtlB, which contributes to septum cleavage and cell wall remodelling (Mesnage et al., 2008).

**Figure 4.**
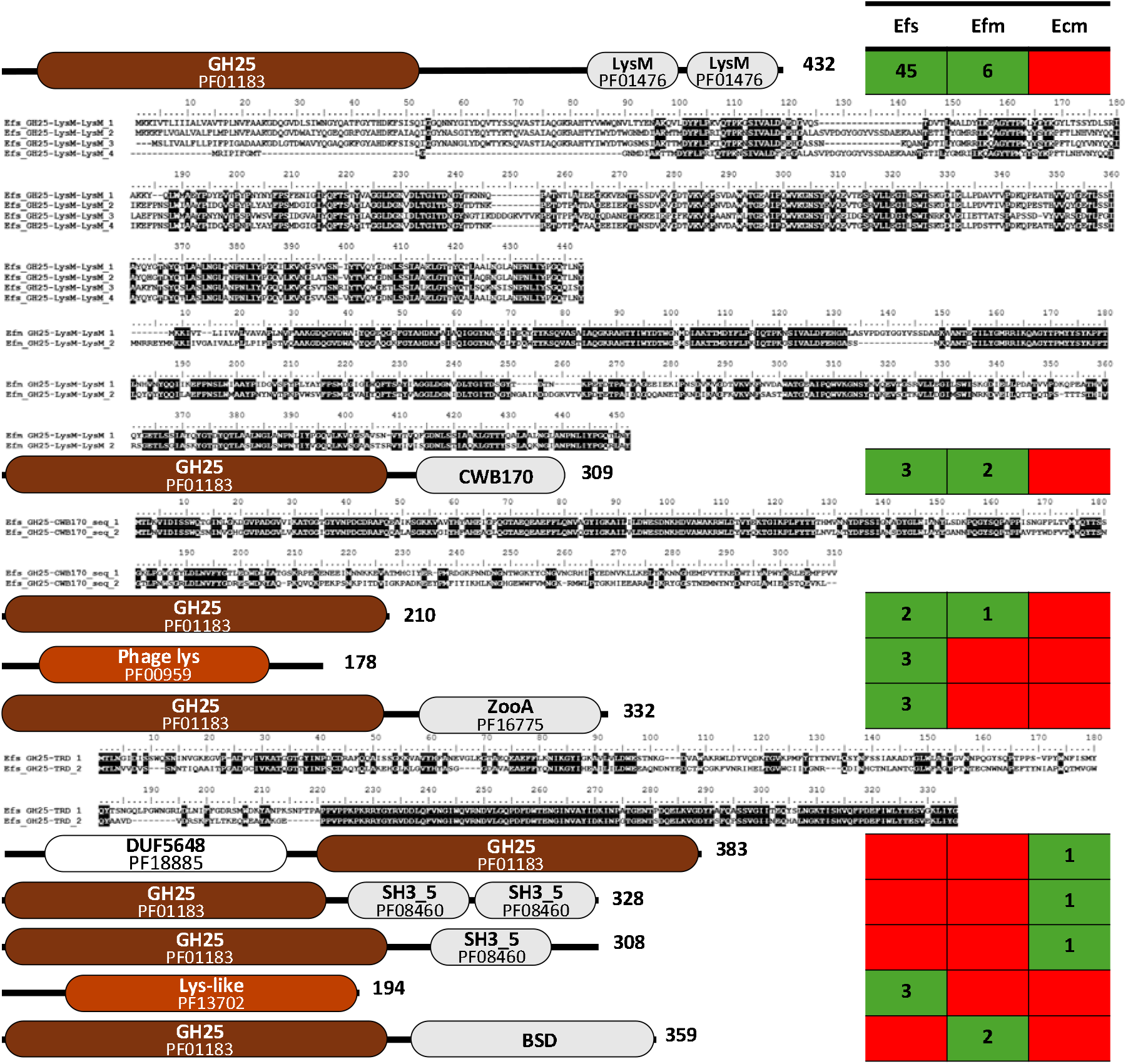
Domain architecture of endolysins containing a muramidase CD. Domain organisations of multiple muramidase containing endolysins in combination with various CWBDs. Boxes on the right-hand side show the number of sequence clusters (95% identity) in each species with a given domain architecture. Further clustering at 75% identity led to the identification of divergent groups of sequences for several muramidase endolysins.

Muramidase CDs were observed in association with several CWBDs: LysM (PF01476), SH3 (PF08240), and ZooA (PF16775), as well as the BSD and CW170 domains. Additionally, a single instance of a GH25 catalytic domain fused to a C-terminal DUF5648 domain was identified in *Ecm*. GH25 domains were only associated with SH3 CWBDs in *Ecm*, in contrast to *Efm* and *Efs* which mostly contain LysM-associated GH25 domains. Domain architectures in *Efs* and *Efm* exhibited greater diversity, each incorporating three distinct identified CWBDs: LysM, ZooA and SH3 (*Efs)*, LysM, CWB170 and BSD (*Efm*).

Structural comparisons of representative muramidase domains again demonstrated high fold conservation, with pairwise structural alignments yielding TM-scores approaching 0.7, indicative of strong structural similarity despite sequence divergence. This structural conservation is consistent with the canonical lysozyme-like GH25 fold which comprises a predominantly α-helical architecture surrounding a conserved catalytic glutamate residue. Sequence diversity was again observed with three of the domain organisations including GH25-LysM-LysM (4 *Efs* and 2 *Efm* clusters), GH25-CWB170 (2 *Efs* clusters) and GH25-ZooA (2 *Efs* clusters) (Supp. File 2).

### Endolysins with multiple catalytic domains

All species studied encoded endolysins with multi-catalytic domain architectures. These included a glycosyl hydrolase domain (GH73 and lysozyme-like) and one or two peptidase domains (M23, NlpC/P60 or CHAP), and a single example of an amidase-like domain. Only one of these multi-catalytic lysins was found to also contain two SH3 CWBDs as well as an unknown C-terminal domain that we called CTD2 (Fig. 5). This architectural complexity is consistent with prior observations that modularity enhances substrate recognition, catalytic efficiency, and host specificity among Gram-positive-targeting lysins (Becker et al., 2016). Interestingly some of the CDs present in these multi-domain endolysins (GH73 and M23) were never detected as standalone CDs. The most common architecture was a GH73 fused to a C-terminal M23 CD with no CWBDs and was found exclusively in *Efm*. Other architectures were seen only once or twice. The 13 GH73-M23 lysins identified in *Efm* included five sequence clusters displaying less than 75% sequence similarity (Supp. File 2), which mainly came from the predicted N-terminal intrinsically disordered region; limited sequence variability was also detected in both CDs.

**Figure 5.**
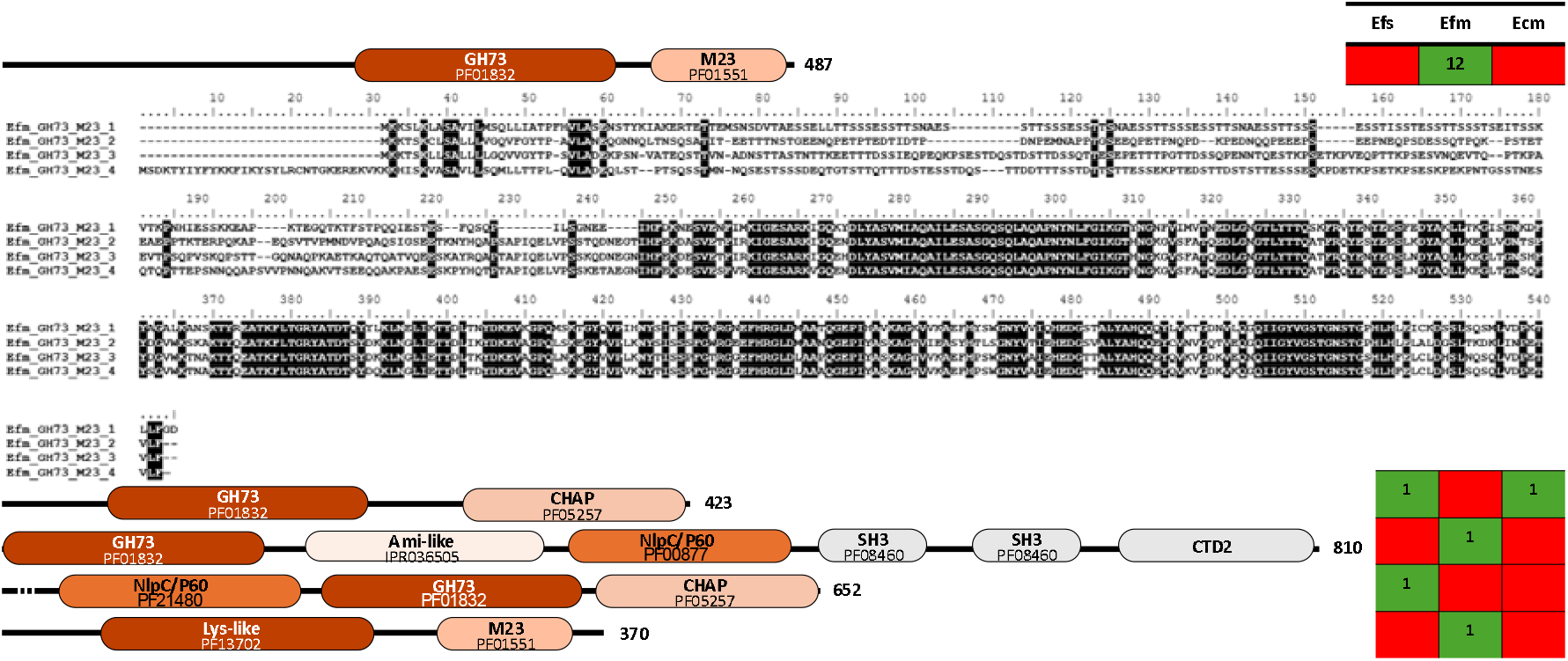
Domain architecture of endolysins with multiple catalytic domains. Domain organisations of various endolysins with multiple catalytic domains are shown. Boxes on the right-hand side show the number of sequence clusters (95% identity) in each species with a given domain architecture. An arbitrary cutoff of 75% identity led to the identification of divergent groups of sequences for several endolysins with multiple CDs.

### In silico characterisation of putative novel cell wall binding domains conserved across species

The endolysins identified in this study contained several well-characterised CWBDs previously described in the literature (Mesnage et al., 2014, Gonzalez-Delgado et al., 2020). In addition to these established domains, we identified three new domains that were present across the three species analysed and which, to our knowledge, have not been previously described. We have designated these novel modules BSD, CTD1, and CTD2. BSD is 135 amino acids in length (Fig. 6A), and structural predictions indicate that the domain adopts a compact fold consisting of two antiparallel β-sheets arranged in a sandwich-like configuration, with a single α-helix positioned at the C-terminal end of the domain (Fig. 6B). Despite limited sequence identity (Fig. 6A), the BSD fold is predicted to be highly conserved (Fig. 6B). CTD1 is 117 amino acids in length and is made up of mainly alpha helices. CTD2 on the other hand, is 122 amino acids in length and is composed of five antiparallel β-strands and two α-helices, one positioned in the middle of the structure and one at the C-terminal end of the protein. These structures seem to form stable scaffolds consistent with a potential role in cell wall recognition and binding.

**Figure 6.**
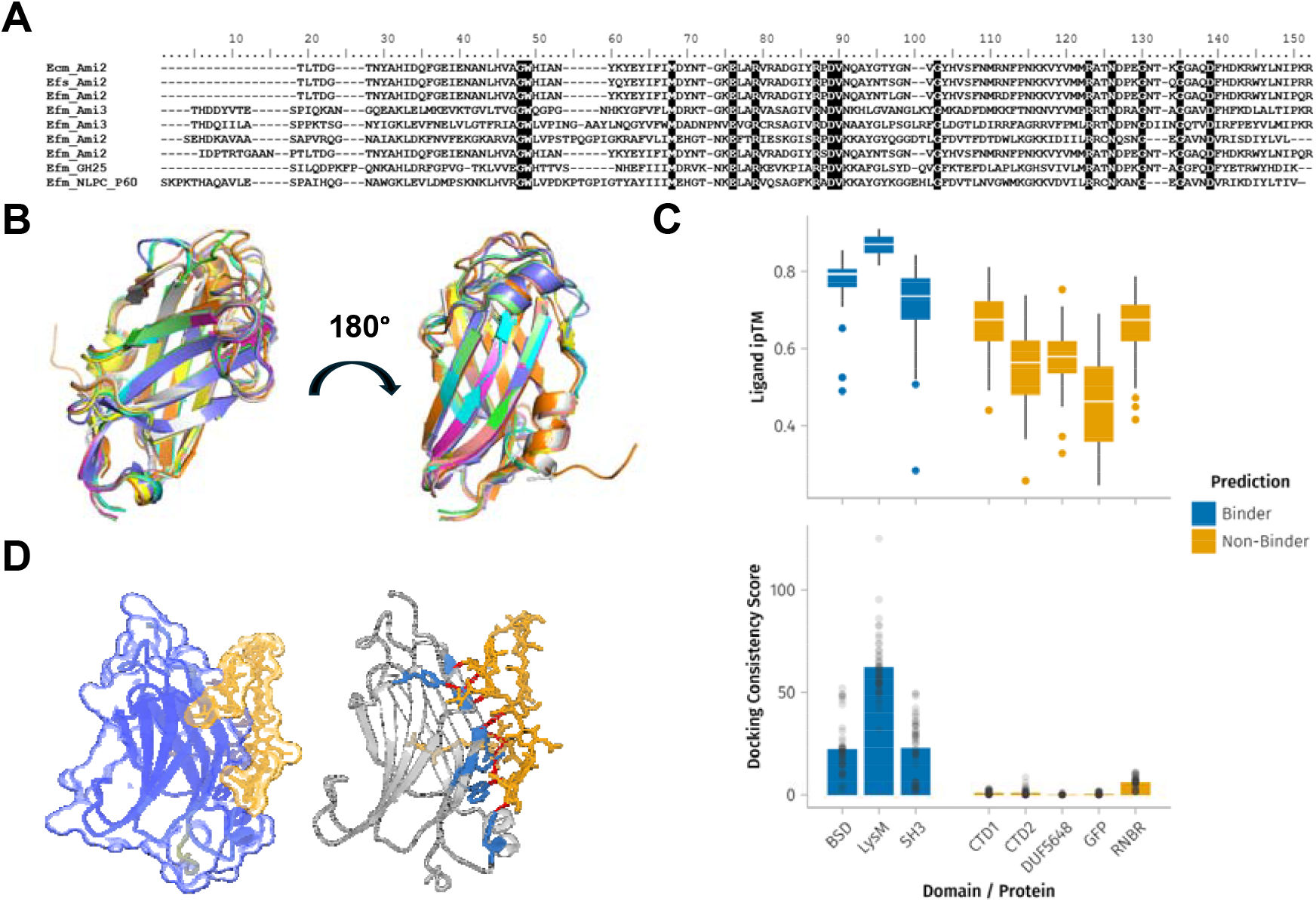
BSD is predicted to be a novel cell wall binding domain. **A**, Alignment of amino acid sequences encoding the BSD. **B**, Structural alignment of AlphaFold predictions. **C**, A box plot quantifying the ligand ipTM scores of 25 *Efm* and *Efs* muropeptide dockings for various domains / proteins, and the custom-calculated docking consistency scores for each of those 25 predictions. The results have been visually grouped into predicted binders and predicted non-binders by comparing ipTM and docking consistency scores to known binders (LysM, SH3) and random proteins (GFP, RNBR). **D**, Docking of *Efs* muropeptide (gold) into the binding pocket of BSD (blue), including any predicted hydrogen bonding.

Notably, BSD was identified in endolysins harbouring distinct catalytic domains, including amidase (Ami_2/Ami_3), endopeptidase (NlpCP60) and muramidase (GH25) modules. This distribution across functionally different enzymatic classes suggests that the domain may act as a modular binding component capable of associating with diverse catalytic architectures. CTD1 was found in association with both amidase (Ami_2) and endopeptidase (NlpC_P60) domains, but CTD2 was only present in a single lysin, being found in a complex with three catalytic domains and two additional CWBDs (both SH3 domains).

To determine if these domains could be novel CWBDs, muropeptide ligands from both *Efs* and *Efm* were docked into them *in silico*. For each docking, the ligand ipTM, a measure of docking confidence, was extracted and plotted in Fig. 6C (Top). Dockings with BSD showed similar ipTM scores to known CWBDs (like LysM and SH3). CTD1 also showed high ipTM scores close to that of BSD indicating this too could be a cell binding domain. CTD2, on the other hand, showed ipTM scores more like non-CWBD controls (RNBR and GFP) suggesting it may not be involved in binding.

Despite displaying similar ipTM scores to known binders and BSD, the ipTM score of CTD1 was also close to that of barnase (RNBR, a ribonuclease included as a negative control). Visualization of the predicted structures revealed that, despite the high ipTM score, the docking position of muropeptides into CTD1 was inconsistent between models. To capture this ligand-pose inconsistency, we developed a custom “docking consistency” score (see Methods). We found this metric better captured the binding pocket and pose consistency that is indicative of a real ligand interaction and more cleanly separated our binder and non-binder controls. Docking consistency scores for our highest confidence binders (BSD, LysM, and SH3 based on ipTM scores) were all high, indicating that the muropeptides were oriented consistently across the 25 models generated by Boltz (Fig, 6C; Bottom). In contrast, RNBR and CTD1 showed low scores, indicating an inconsistency in muropeptide orientation across models. As a result of this inconsistency, we were unable to confidently predict CTD1 to be a cell binding domain.

Muropeptides from *Efs* (gmgm-AQK[AA]AA∼gm-AQK[AA]AA∼g) and *Efm* (gmgm-AQK[D]AA∼gm-AQK[D]AA∼g) fit neatly into defined grooves on the surface of the BSD (Fig. 6D; 3D models in Supp File. 3 ‘BSD Visualisations/’) and formed ∼10 hydrogen bonds to the protein.

Interestingly, almost all those hydrogen bonds were to BSD’s peptide backbone, not to its amino acid sidechains. This may help to explain the relative lack of conserved amino acids in BSD sequences, as it suggests only the conserved backbone fold is required for muropeptide binding. Collectively, these findings support the classification of BSD as a structurally conserved and potentially functionally significant CWBD in endolysins. It should be noted that, despite the low ipTM and docking consistency scores for both CTD1 and CTD2, their positioning relative to catalytic domains (being found at the C-terminus) is similar to that of the other CWBDs identified in this study. This suggests the domains could still be involved in cell wall binding, though perhaps via non-peptidoglycan component of the cell wall or untested peptidoglycan substrate.

## Concluding remarks

This study significantly expands the known set of phage endolysins that could potentially be used as enzybiotics for the treatment of enterococcal infections. Data mining of prophage genomes revealed 33 unique architectures of enterococcal phage endolysins, approximately twice as many as previously reported from analyses of lytic phages. Our enumeration of the sequence diversity within each domain architecture serves as a valuable resource from which recombinant proteins can be expressed and evaluated as antimicrobials. The high□throughput expression and screening of these recombinant endolysins, and potentially the creation of new, domain-shuffled endolysins represent powerful strategies for the discovery and development of new, phage□derived therapeutics.

## Acknowledgements

Work in the SM lab was funded by BBSRC (grant BB/W013800/1). FOD is the recipient of a White Rose Doctoral Training Programme iCASE studentship (grant BB/T007222/1), BJR is the recipient of a NERC PhD iCASE studentship (NERC ACCE NE/S00713X/1). AK is funded by a PhD studentship from the Medical Research Council (grant MR/W007002/1).

## References

Abramson, J., Adler, J., Dunger, J., Evans, R., Green, T., Pritzel, A., Ronneberger, O., Willmore, L., Ballard, A. J., Bambrick, J., Bodenstein, S. W., Evans, D. A., Hung, C. C., O’Neill, M., Reiman, D., Tunyasuvunakool, K., Wu, Z., ŽEmgulyte, A., Arvaniti, E., Beattie, C., Bertolli, O., Bridgland, A., Cherepanov, A., Congreve, M., Cowen-Rivers, A. I., Cowie, A., Figurnov, M., Fuchs, F. B., Gladman, H., Jain, R., Khan, Y. A., Low, C. M. R., Perlin, K., Potapenko, A., Savy, P., Singh, S., Stecula, A., Thillaisundaram, A., Tong, C., Yakneen, S., Zhong, E. D., Zielinski, M., ŽíDek, A., Bapst, V., Kohli, P., Jaderberg, M., Hassabis, D. & Jumper, J. M. 2024. Accurate structure prediction of biomolecular interactions with AlphaFold 3. Nature, 630, 493–500.

AlamáN-ZáRate, M. G., Rady, B. J., Ledermann, R., Shephard, N., Evans, C. A., Dickman, M. J., Turner, R. D., Rifflet, A., Patel, A. V., Gomperts Boneca, I., Poole, P. S., Bern, M. & Mesnage, S. 2025. A software tool and strategy for peptidoglycomics, the high-resolution analysis of bacterial peptidoglycans via LC-MS/MS. Commun Chem, 8, 91.

Alrafaie, A. M. & Stafford, G. P. 2023. Enterococcal bacteriophage: A survey of the tail associated lysin landscape. Virus Res, 327, 199073.

Balaban, C. L., SuáRez, C. A., Boncompain, C. A., Peressutti-Bacci, N., Ceccarelli, E. A. & Morbidoni, H. R. 2022. Evaluation of factors influencing expression and extraction of recombinant bacteriophage endolysins in Escherichia coli. Microb Cell Fact, 21, 40.

Becker, S. C., Roach, D. R., Chauhan, V. S., Shen, Y., Foster-Frey, J., Powell, A. M., Bauchan, G., Lease, R. A., Mohammadi, H., Harty, W. J., Simmons, C., Schmelcher, M., Camp, M., Dong, S., Baker, J. R., Sheen, T. R., Doran, K. S., Pritchard, D. G., Almeida, R. A., Nelson, D. C., Marriott, I., Lee, J. C. & Donovan, D. M. 2016. Triple-acting Lytic Enzyme Treatment of Drug-Resistant and Intracellular Staphylococcus aureus. Sci Rep, 6, 25063.

Beganovic, M., Luther, M. K., Rice, L. B., Arias, C. A., Rybak, M. J. & Laplante, K. L. 2018. A Review of Combination Antimicrobial Therapy for Enterococcus faecalis Bloodstream Infections and Infective Endocarditis. Clin Infect Dis, 67, 303–309.

Bezanson, J., Edelman, A., Karpinski, S. & Shah, V. B. 2017. Julia: A Fresh Approach to Numerical Computing. SIAM Review, 59, 65–98.

Blum, M., Andreeva, A., Florentino, L. C., Chuguransky, S. R., Grego, T., Hobbs, E., Pinto, B. L., Orr, A., Paysan-Lafosse, T., Ponamareva, I., Salazar, G. A., Bordin, N., Bork, P., Bridge, A., Colwell, L., Gough, J., Haft, D. H., Letunic, I., Llinares-LóPez, F., Marchler-Bauer, A., Meng-Papaxanthos, L., Mi, H., Natale, D. A., Orengo, C. A., Pandurangan, A. P., Piovesan, D., Rivoire, C., Sigrist, C. J. A., Thanki, N., Thibaud-Nissen, F., Thomas, P. D., Tosatto, S. C. E., Wu, C. H. & Bateman, A. 2025. InterPro: the protein sequence classification resource in 2025. Nucleic Acids Res, 53, D444–d456.

Bustamante, N., Iglesias-Bexiga, M., Bernardo-GarcíA, N., Silva-MartíN, N., GarcíA, G., Campanero-Rhodes, M. A., GarcíA, E., UsóN, I., Buey, R. M., GarcíA, P., Hermoso, J. A., Bruix, M. & MenÉNdez, M. 2017. Deciphering how Cpl-7 cell wall-binding repeats recognize the bacterial peptidoglycan. Sci Rep, 7, 16494.

Camargo, A. P., Roux, S., Schulz, F., Babinski, M., Xu, Y., Hu, B., Chain, P. S. G., Nayfach, S. & Kyrpides, N. C. 2024. Identification of mobile genetic elements with geNomad. Nat Biotechnol, 42, 1303–1312.

Chaumeil, P. A., Mussig, A. J., Hugenholtz, P. & Parks, D. H. 2022. GTDB-Tk v2: memory friendly classification with the genome taxonomy database. Bioinformatics, 38, 5315–5316.

Cheng, M., Zhang, Y., Li, X., Liang, J., Hu, L., Gong, P., Zhang, L., Cai, R., Zhang, H., Ge, J., Ji, Y., Guo, Z., Feng, X., Sun, C., Yang, Y., Lei, L., Han, W. & Gu, J. 2017. Endolysin LysEF-P10 shows potential as an alternative treatment strategy for multidrug-resistant Enterococcus faecalis infections. Sci Rep, 7, 10164.

Codelia-Anjum, A., Lerner, L. B., Elterman, D., Zorn, K. C., Bhojani, N. & Chughtai, B. 2023. Enterococcal Urinary Tract Infections: A Review of the Pathogenicity, Epidemiology, and Treatment. Antibiotics (Basel), 12.

Danisch et al. 2021. Makie.jl: Flexible high-performance data visualization for Julia. Journal of Open Source Software, 6(65).

Dedrick, R. M., Smith, B. E., Cristinziano, M., Freeman, K. G., Jacobs-Sera, D., Belessis, Y., Whitney Brown, A., Cohen, K. A., Davidson, R. M., Van Duin, D., Gainey, A., Garcia, C. B., Robert George, C. R., Haidar, G., Ip, W., Iredell, J., Khatami, A., Little, J. S., Malmivaara, K., Mcmullan, B. J., Michalik, D. E., Moscatelli, A., Nick, J. A., Tupayachi Ortiz, M. G., Polenakovik, H. M., Robinson, P. D., Skurnik, M., Solomon, D. A., Soothill, J., Spencer, H., Wark, P., Worth, A., Schooley, R. T., Benson, C. A. & Hatfull, G. F. 2022. Phage Therapy of Mycobacterium Infections: Compassionate Use of Phages in 20 Patients With Drug-Resistant Mycobacterial Disease. Clinical Infectious Diseases, 76, 103–112.

Dolka, B., Chrobak-Chmiel, D., Makrai, L. & Szeleszczuk, P. 2016. Phenotypic and genotypic characterization of Enterococcus cecorum strains associated with infections in poultry. BMC Vet Res, 12, 129.

Donelli, G. & Guaglianone, E. 2004. Emerging role of Enterococcus spp in catheter-related infections: biofilm formation and novel mechanisms of antibiotic resistance. J Vasc Access, 5, 3–9.

Feldwisch, J., Tolmachev, V., Lendel, C., Herne, N., SjÖBerg, A., Larsson, B., Rosik, D., Lindqvist, E., Fant, G., HÖIdÉN-Guthenberg, I., Galli, J., Jonasson, P. & AbrahmsÉN, L. 2010. Design of an optimized scaffold for affibody molecules. J Mol Biol, 398, 232–47.

Fischetti, V. A. 2005. Bacteriophage lytic enzymes: novel anti-infectives. Trends Microbiol, 13, 491–6.

Fu, L., Niu, B., Zhu, Z., Wu, S. & Li, W. 2012. CD-HIT: accelerated for clustering the next-generation sequencing data. Bioinformatics, 28, 3150–2.

Giazitzidou, S. P. A. S. 2018. A SYNTHESIS OF RESEARCH ON READING FLUENCY DEVELOMPENT: STUDY OF EIGHT META-ANALYSES. European Journal of Special Education Research, 3(4).

Gondil, V. S., Harjai, K. & Chhibber, S. 2020. Endolysins as emerging alternative therapeutic agents to counter drug-resistant infections. Int J Antimicrob Agents, 55, 105844.

Gonzalez-Delgado, L. S., Walters-Morgan, H., Salamaga, B., Robertson, A. J., Hounslow, A. M., Jagielska, E., SabałA, I., Williamson, M. P., Lovering, A. L. & Mesnage, S. 2020. Two-site recognition of Staphylococcus aureus peptidoglycan by lysostaphin SH3b. Nat Chem Biol, 16, 24–30.

Greener, J. G., Selvaraj, J. & Ward, B. J. 2020. BioStructures.jl: read, write and manipulate macromolecular structures in Julia. Bioinformatics, 36, 4206–4207.

Hatfull, G. F., Dedrick, R. M. & Schooley, R. T. 2022. Phage Therapy for Antibiotic-Resistant Bacterial Infections. Annu Rev Med, 73, 197–211.

HEALTH, U. D. O. & SERVICES, H. 2019. Antibiotic resistance threats in the United States, 2019. (No Title).

Hollenbeck, B. L. & Rice, L. B. 2012. Intrinsic and acquired resistance mechanisms in enterococcus. Virulence, 3, 421–33.

Kieft, K., Zhou, Z. & Anantharaman, K. 2020. VIBRANT: automated recovery, annotation and curation of microbial viruses, and evaluation of viral community function from genomic sequences. Microbiome, 8, 90.

Kim, S., Jin, J. S., Choi, Y. J. & Kim, J. 2020. LysSAP26, a New Recombinant Phage Endolysin with a Broad Spectrum Antibacterial Activity. Viruses, 12.

Kovalskaya, N. Y., Herndon, E. E., Foster-Frey, J. A., Donovan, D. M. & Hammond, R. W. 2019. Antimicrobial activity of bacteriophage derived triple fusion protein against Staphylococcus aureus. AIMS Microbiol, 5, 158–175.

Kristich, C. J., Rice, L. B. & Arias, C. A. 2014. Enterococcal infection—treatment and antibiotic resistance.

Lai, A. C., Tran, S. & Simmonds, R. S. 2002. Functional characterization of domains found within a lytic enzyme produced by Streptococcus equi subsp. zooepidemicus. FEMS Microbiol Lett, 215, 133–8.

Laurentie, J., Mourand, G., Grippon, P., Furlan, S., Chauvin, C., Jouy, E., Serror, P. & Kempf, I. 2023. Determination of Epidemiological Cutoff Values for Antimicrobial Resistance of Enterococcus cecorum. J Clin Microbiol, 61, e0144522.

Liu, H., Hu, Z., Li, M., Yang, Y., Lu, S. & Rao, X. 2023. Therapeutic potential of bacteriophage endolysins for infections caused by Gram-positive bacteria. J Biomed Sci, 30, 29.

Loeffler, J. M., Nelson, D. & Fischetti, V. A. 2001. Rapid killing of Streptococcus pneumoniae with a bacteriophage cell wall hydrolase. Science, 294, 2170–2.

Loessner, M. J. 2005. Bacteriophage endolysins--current state of research and applications. Curr Opin Microbiol, 8, 480–7.

Mesnage, S., Chau, F., Dubost, L. & Arthur, M. 2008. Role of N-acetylglucosaminidase and N-acetylmuramidase activities in Enterococcus faecalis peptidoglycan metabolism. J Biol Chem, 283, 19845–53.

Mesnage, S., Dellarole, M., Baxter, N. J., Rouget, J. B., Dimitrov, J. D., Wang, N., Fujimoto, Y., Hounslow, A. M., Lacroix-Desmazes, S., Fukase, K., Foster, S. J. & Williamson, M. P. 2014. Molecular basis for bacterial peptidoglycan recognition by LysM domains. Nat Commun, 5, 4269.

Mitkowski, P., Jagielska, E. & SabałA, I. 2024. Engineering of chimeric enzymes with expanded tolerance to ionic strength. Microbiol Spectr, 12, e0354623.

Naddaf, M. 2024. 40 million deaths by 2050: toll of drug-resistant infections to rise by 70. Nature, 633, 747–748.

Nelson, D., Loomis, L. & Fischetti, V. A. 2001. Prevention and elimination of upper respiratory colonization of mice by group A streptococci by using a bacteriophage lytic enzyme. Proc Natl Acad Sci U S A, 98, 4107–12.

O’Dea, F., Dupuis, A., Brown, J., Edwards, J., Kay, S., Stafford, G. P. & Mesnage, S. 2025. Phenotypic and genomic analysis of the emerging poultry pathogen Enterococcus cecorum in UK isolates. Microbial Genomics, 11.

Onallah, H., Hazan, R., Nir-Paz, R., Brownstein, M. J., Fackler, J. R., Horne, B., Hopkins, R., Basu, S., Yerushalmy, O., Alkalay-Oren, S., Braunstein, R., Rimon, A., Gelman, D., Khalifa, L., Adler, K., Abdalrhman, M., Gelman, S., Katvan, E., Coppenhagen-Glazer, S., Moses, A., Oster, Y., Dekel, M., Ben-Ami, R., Khoury, A., Kedar, D. J., Meijer, S. E., Ashkenazi, I., Bishouty, N., Yahav, D., Shostak, E., Livni, G., Paul, M., Gross, M., Ormianer, M., Aslam, S., Ritter, M., Urish, K. L., La Hoz, R. M., Khatami, A., Britton, P. N., Lin, R. C. Y., Iredell, J. R., Petrovic-Fabijan, A., Lynch, S., Tamma, P. D., Yamshchikov, A., Lesho, E., Morales, M., Werzen, A. & Saharia, K. 2023. Refractory Pseudomonas aeruginosa infections treated with phage PASA16: A compassionate use case series. Med, 4, 600-611.e4.

ORGANIZATION, W. H. 2014. Antimicrobial resistance: global report on surveillance, World Health Organization.

Parks, D. H., Imelfort, M., Skennerton, C. T., Hugenholtz, P. & Tyson, G. W. 2015. CheckM: assessing the quality of microbial genomes recovered from isolates, single cells, and metagenomes. Genome Res, 25, 1043–55.

Passaro, S., Corso, G., Wohlwend, J., Reveiz, M., Thaler, S., Somnath, V. R., Getz, N., Portnoi, T., Roy, J., Stark, H., Kwabi-Addo, D., Beaini, D., Jaakkola, T. & Barzilay, R. 2025. Boltz-2: Towards Accurate and Efficient Binding Affinity Prediction. bioRxiv.

Pettersen, E. F., Goddard, T. D., Huang, C. C., Meng, E. C., Couch, G. S., Croll, T. I., Morris, J. H. & Ferrin, T. E. 2021. UCSF ChimeraX: Structure visualization for researchers, educators, and developers. Protein Sci, 30, 70–82.

RodríGuez-Rubio, L., GutiÉRrez, D., Donovan, D. M., MartíNez, B., RodríGuez, A. & GarcíA, P. 2016. Phage lytic proteins: biotechnological applications beyond clinical antimicrobials. Crit Rev Biotechnol, 36, 542–52.

Schwartzman, J. A., Lebreton, F., Salamzade, R., Martin, M. J., Schaufler, K., Urhan, A., Abeel, T., Camargo, I., Sgardioli, B. F., Prichula, J., Frazzon, A. P. G., Van Tyne, D., Treinish, G., Innis, C. J., Wagenaar, J. A., Whipple, R. M., Manson, A. L., Earl, A. M. & Gilmore, M. S. 2023. Global diversity of enterococci and description of 18 novel species. bioRxiv.

Sievers, F., Wilm, A., Dineen, D., Gibson, T. J., Karplus, K., Li, W., Lopez, R., Mcwilliam, H., Remmert, M., SÖDing, J., Thompson, J. D. & Higgins, D. G. 2011. Fast, scalable generation of high-quality protein multiple sequence alignments using Clustal Omega. Mol Syst Biol, 7, 539.

SirÉN, K., Millard, A., Petersen, B., Gilbert, M. T. P., Clokie, M. R. J. & Sicheritz-PontÉN, T. 2021. Rapid discovery of novel prophages using biological feature engineering and machine learning. NAR Genom Bioinform, 3, qaa109.

Souillard, R., Laurentie, J., Kempf, I., LE CaèR, V., Le Bouquin, S., Serror, P. & Allain, V. 2022. Increasing incidence of Enterococcus-associated diseases in poultry in France over the past 15 years. Vet Microbiol, 269, 109426.

Teklemariam, A. D., Al-Hindi, R. R., Qadri, I., Alharbi, M. G., Ramadan, W. S., Ayubu, J., Al-Hejin, A. M., Hakim, R. F., Hakim, F. F., Hakim, R. F., Alseraihi, L. I., Alamri, T. & Harakeh, S. 2023. The Battle between Bacteria and Bacteriophages: A Conundrum to Their Immune System. Antibiotics (Basel), 12.

Terzian, P., Olo Ndela, E., Galiez, C., Lossouarn, J., Pérez Bucio, R.E., Mom, R., Toussaint, A., Petit, M. A. & Enault, F. 2021. PHROG: families of prokaryotic virus proteins clustered using remote homology. NAR Genom Bioinform, 3, qab067.

Uchiyama, J., Takemura, I., Hayashi, I., Matsuzaki, S., Satoh, M., Ujihara, T., Murakami, M., Imajoh, M., Sugai, M. & Daibata, M. 2011. Characterization of lytic enzyme open reading frame 9 (ORF9) derived from Enterococcus faecalis bacteriophage phiEF24C. Appl Environ Microbiol, 77, 580–5.

Van Kempen, M., Kim, S. S., Tumescheit, C., Mirdita, M., Lee, J., Gilchrist, C. L. M., SÖDing, J. & Steinegger, M. 2024. Fast and accurate protein structure search with Foldseek. Nat Biotechnol, 42, 243–246.

Viglasky, J., Piknova, M. & Pristas, P. 2023. Gene and domain shuffling in lytic cassettes of Enterococcus spp. bacteriophages. 3 Biotech, 13, 388.

Wang, C., Zhao, J., Lin, Y., Lwin, S. Z. C., El-Telbany, M., Masuda, Y., Honjoh, K. I. & Miyamoto, T. 2024. Characterization of Two Novel Endolysins from Bacteriophage PEF1 and Evaluation of Their Combined Effects on the Control of Enterococcus faecalis Planktonic and Biofilm Cells. Antibiotics (Basel), 13.

Wang, J., Chitsaz, F., Derbyshire, M. K., Gonzales, N. R., Gwadz, M., Lu, S., Marchler, G. H., Song, J. S., Thanki, N., Yamashita, R. A., Yang, M., Zhang, D., Zheng, C., Lanczycki, C. J. & Marchler-Bauer, A. 2023. The conserved domain database in 2023. Nucleic Acids Res, 51, D384–d388.

Xu, X., Zhang, D., Zhou, B., Zhen, X. & Ouyang, S. 2021. Structural and biochemical analyses of the tetrameric cell binding domain of Lys170 from enterococcal phage F170/08. Eur Biophys J, 50, 721–729.

Yoong, P., Schuch, R., Nelson, D. & Fischetti, V. A. 2004. Identification of a broadly active phage lytic enzyme with lethal activity against antibiotic-resistant Enterococcus faecalis and Enterococcus faecium. J Bacteriol, 186, 4808–12.

Zhou B, Zhen X, Zhou H, Zhao F, Fan C, Perčulija V, Tong Y, Mi Z, Ouyang S. Structural and functional insights into a novel two-component endolysin encoded by a single gene in Enterococcus faecalis phage. PLoS Pathog. 2020 Mar 16;16(3):e1008394. doi: 10.1371/journal.ppat.1008394. PMID: 32176738; PMCID: PMC7098653

